# PolyQ-independent toxicity associated with novel translational products from CAG repeat expansions

**DOI:** 10.1101/2019.12.20.884254

**Authors:** Paige D. Rudich, Simon Watkins, Todd Lamitina

## Abstract

Expanded CAG nucleotide repeats are the underlying genetic cause of at least 14 incurable diseases, including Huntington’s disease (HD). The toxicity associated with many CAG repeat expansions is thought to be due to the translation of the CAG repeat to create a polyQ protein, which forms toxic oligomers and aggregates. However, recent studies show that HD CAG repeats undergo a non-canonical form of translation called <u>R</u>epeat-<u>a</u>ssociated <u>n</u>on-AUG dependent (RAN) translation. RAN translation of the CAG sense and CUG anti-sense RNAs produces six distinct repeat peptides: polyalanine (polyAla, from both CAG and CUG repeats), polyserine (polySer), polyleucine (polyLeu), polycysteine (polyCys), and polyglutamine (polyGln). The toxic potential of individual CAG-derived RAN polypeptides is not well understood. We developed pure *C. elegans* protein models for each CAG RAN polypeptide using codon-varied expression constructs that preserve RAN protein sequence but eliminate repetitive CAG/CUG RNA. While all RAN polypeptides formed aggregates, only polyLeu was consistently toxic across multiple cell types. In GABAergic neurons, which exhibit significant neurodegeneration in HD patients, codon-varied (Leu)_38_, but not (Gln)_38_, caused substantial neurodegeneration and motility defects. Our studies provide the first *in vivo* evaluation of CAG-derived RAN polypeptides and suggest that polyQ-independent mechanisms, such as RAN-translated polyLeu peptides, may have a significant pathological role in CAG repeat expansion disorders.

## Introduction

DNA repeat expansions are the genetic cause of >30 different diseases, most of which affect the nervous system and are currently incurable [1]. Of these, CAG/CTG repeat expansions are associated with at least 14 neurodegenerative diseases. CAG/CTG repeat expansions can occur in either protein-coding exons or untranslated regions. Diseases caused by exonic CAG repeat expansions are often referred to as ‘polyQ’ diseases, because the expanded CAG repeat is in-frame with the encoded protein and leads to the translation of a series of glutamine (Q) residues. Expanded polyQ proteins form large protein aggregates that sequester many other cellular proteins and undergo significant post-translational modifications, such as ubiquitination [2–5]. While these polyQ aggregates were initially thought to be toxic, other studies suggest that large aggregates may be protective and that smaller oligomers containing polyQ proteins are the toxic entity [6], leading to mitochondrial disruption, alterations in the protein folding landscape, and impaired autophagy [7–9]. Despite the dogma that expanded polyQ proteins are the molecular cause of toxicity in exonic CAG repeat expansions, significant data suggests that polyQ oligomers or aggregates may not always be associated with toxicity. For example, in adult Huntington’s disease (HD) post-mortem tissues, some white matter regions of the caudate and putamen exhibit significant neurodegeneration but lack detectable polyQ aggregates [10]. Similarly, in juvenile HD (a more aggressive form of HD associated with higher numbers of CAG repeats), the cerebellum undergoes significant neurodegeneration, but lacks polyQ aggregates in post-mortem tissues [10]. In a recent study, patient genetic data shows that uninterrupted CAG repeat length determines the age of disease onset in HD through a mechanism separate from its glutamine-coding property [11, 12]. These observations suggest that polyQ-independent toxicity mechanisms may play a role in CAG repeat expansion disorders like HD.

One possible polyQ-independent mechanism involves a recently discovered type of protein translation, <u>r</u>epeat-<u>a</u>ssociated <u>n</u>on-AUG dependent (RAN) translation [13]. RAN translation uses as a substrate the long, G/C-rich RNA repeats that are typically found in repeat expansion diseases. As RAN translation does not require a start codon, these G/C-rich RNAs are translated in each reading frame, through a mechanism that is still under investigation [14–16]. Additionally, repeat-expansion DNA commonly produces an antisense RNA, which is also G/C rich and thus can also undergo RAN translation [17, 18]. RAN translation occurs in both exonic CAG expansion diseases (HD [10] and spinocerebellar ataxia type 8 (SCA8) [19]) and an untranslated CTG expansion disease (the 3’UTR CTG expansion disease, myotonic dystrophy type 1 (DM1) [17]). In both adult and juvenile HD patients, HD RAN products are present in brain regions lacking polyQ that are still undergoing apoptosis [10]. Thus, RAN-translated peptides could be a polyQ-independent mechanism that contributes to the pathology of many or all CAG/CTG repeat expansion diseases.

Identifying which of the CAG RAN polypeptides are pathogenic and their mechanism(s) of toxicity is necessary to understand the toxic potential of CAG/CTG-derived RAN polypeptides. Canonical translation of CAG RNA repeats produces polyglutamine (polyGln). However, the same RNA also undergoes RAN translation to produce polyserine (polySer), and polyalanine (polyAla) [10]. RAN translation of antisense CUG RNA repeats produces polyleucine (polyLeu), polycysteine (polyCys), and polyalanine (polyAla) [10]. All four RAN polypeptides and polyQ are weakly toxic at 90 repeats and at least two peptides (polySer and polyGln) form protein aggregates when overexpressed in cultured cells [10]. Whether or not more disease-relevant lengths of these peptides are also toxic and form protein aggregates is not known.

To begin addressing these questions, we created codon-varied GFP-tagged HD RAN homopolymers at both disease-relevant lengths (38 repeats) and highly expanded lengths (90 repeats) using previously described approaches [20]. We expressed these peptides in multiple cellular settings in *C. elegans*. We found that each RAN peptide formed protein aggregates. However, only polyLeu was toxic in all cellular settings. PolyLeu displayed length-associated toxicity and caused significant neurotoxicity *in vivo*. Notably, neurotoxicity was not observed with codon-varied polyGln, suggesting that these two polypeptides act through distinct pathways. Identifying the cellular processes involved in polyLeu toxicity may provide new biomarkers and therapeutic targets for HD and other CAG repeat expansion disorders.

## Results

### Development of a *C. elegans* HD RAN Model

To evaluate the toxicity of individual CAG-encoded RAN polypeptides, we created *C. elegans* models for each of the possible CAG RAN products using a previously described codon-variation strategy. This approach maintains the amino acid repeat of the RAN polypeptide but removes the CAG nucleotide repeat [20]. The codon-varied constructs were also designed to minimize computationally predicted hairpin structures in the resulting RNA, as hairpin structures are thought to be required for RAN translation and could lead to the production of non-relevant RAN products [21]. The codon-varied RAN polypeptides were expressed through canonical AUG-initiated translation and studied at either 38 repeats or 90 repeats with a C-terminal GFP tag (Figure 1A). The 38-repeat peptides mimic the minimum CAG repeat threshold for developing HD. The 90-repeat peptides mimic the length investigated in a previous study [22]. The modeled RAN polypeptides lacked genetic context (i.e. no flanking human HTT sequence). We chose not to include such genetic context because the unique reading frames of each RAN product means that no two RAN polypeptides share the same flanking sequence. Including the genetic contexts of the RAN polypeptides would limit our ability to assign phenotypes to specific homopolymeric peptides rather than flanking sequences. In addition to the individual RAN polypeptides, we also created an AUG-initiated pure CAG repeat in the polyglutamine reading frame, which mimics previous CAG/polyQ models in *C. elegans* [23–27]. This pure CAG repeat could exhibit RNA toxicity, RAN polypeptide toxicity, or a combination of both. For clarity, we will refer to the CAG-encoded polyglutamine as “polyQ” and the codon-varied polyglutamine as “polyGln”. We expressed each construct in both GABAergic neurons and muscle cells using cell-type-specific promoters.

**Figure 1.**
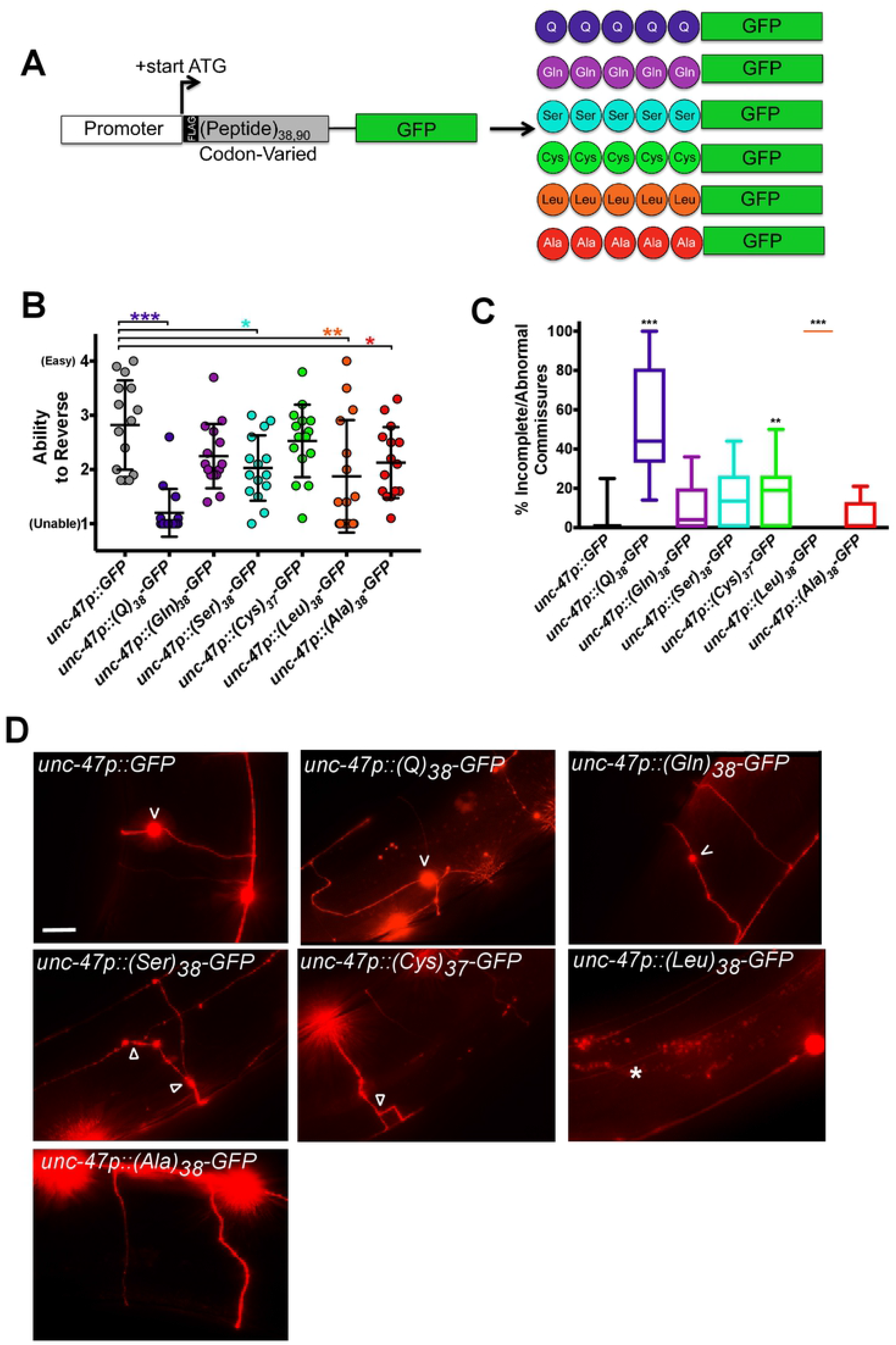
Codon-varied polyLeu, but not polyGln, causes phenotypic and morphological defects in GABAergic neurons. (A) Molecular strategy for expression of codon-varied polypeptide repeats in *C. elegans*. (B) Quantification of the reversal ability of transgenic animals expressing the indicated polypeptide under the GABAergic neuron specific *unc-47* promoter. N = 15 animals/genotype. Each symbol represents one animal, the horizontal line is the mean, and the bars define the standard deviation. *P<0.05, **P<0.01, ***P<0.001 versus GFP control (one-way non-parametric ANOVA with Dunn’s post hoc test). (C) Percentage of incomplete or abnormal commissures / total number of commissures. N = 11-20 animals/genotype. The data is expressed in a box and whisker plot where the whiskers define min and max values. **P<0.01, ***P<0.001 versus GFP control (one-way nonparametric ANOVA with Dunn’s post hoc test). (D) Representative images of *unc-47*+ motor neurons in animals expressing the indicated codon-varied transgene. ‘V’ points to neuronal blebs, arrowhead points to branching, and the asterisk indicates a commissure which fails to reach the dorsal side. Scale bar = 10 μm.

### GABAergic Neuron Model of HD RAN Polypeptides

Neurodegeneration in HD initiates in the striatum and most strongly degrades the GABAergic neuronal population [28]. HD RAN polypeptides are found in degenerating regions of the striatum which lack polyQ, suggesting they contribute to neurodegeneration [22]. We expressed RAN polypeptides in GABAergic motor neurons using the *unc-47* promoter [29] and quantified GABAergic neuron commissures, as they are commonly used to measure neurodegeneration because they provide single axon resolution [30, 31]. As a measurement of GABAergic neuron function, we evaluated directional reversals since loss of GABAergic neurons significantly decreases the ability of *C. elegans* to reverse direction [29]. Quantifying the number, morphology, and function of GABAergic neurons is a common measure of the neurotoxic potential of proteins expressed in *C. elegans* [31–34].

(Q)_38_-GFP worms, which expressed CAG-encoded polyglutamine (and possibly other RAN polypeptides), strongly inhibited the ability of the worms to reverse when expressed in GABAergic neurons (Figure 1B). Surprisingly, (Gln)_38_-GFP worms expressing a codon-varied polyglutamine were comparable to control animals expressing GFP alone. In contrast, (Ser)_38_-GFP, (Leu)_38_-GFP, and (Ala)_38_-GFP each caused reversal defects (Figure 1B). No defects were observed for (Cys)_37_-GFP.

To test if the functional defects were also associated with cellular damage, we visualized neuronal commissures using a GABAergic-neuron RFP marker [31, 32, 35, 36]. Consistent with our functional data, animals expressing (Gln)_38_-GFP exhibited no defects in commissure structure (Figure 1C, D). Animals expressing (Leu)_38_-GFP exhibited highly penetrant commissure defects, although the neuron somas were intact. Where they were visible, the GABAergic neurons in (Leu)_38_-GFP worms traversed the length of the animals instead of the width, suggesting defects in neuron stability and/or axon guidance (Figure 1C,D). (Q)_38_-GFP worms also had abnormal commissures, with roughly half of the commissures in each worm being abnormal or incomplete. Although (Cys)_37_-GFP did not cause a significant functional defect (Figure 1B), 19% of the commissures in (Cys)_37_-GFP worms were abnormal (Figure 1C,D). (Ser)_38_-GFP did not cause a significant defect in commissure morphology (Figure 1C). These data show that the expression of (Leu)_38_-GFP, but not other RAN polypeptides, is sufficient to cause both structural and functional defects in GABAergic neurons. They also suggest that factors other than polyglutamine contribute to the toxicity of (Q)_38_-GFP, as a codon-varied (Gln)_38_-GFP did not exhibit functional or morphological toxicity.

### Muscle Expression of HD RAN Polypeptides

*C. elegans* GABAergic neurons are anatomically small (cell bodies 1-2 micron) and are inaccessible to some genetic approaches, such as RNAi-mediated gene knockdown [37]. To gain more insights into RAN polypeptide cell biology, as well as to generate a platform for future RNAi and forward mutagenesis-based genetic suppressor screening, we expressed the HD RAN polypeptides in muscle cells using the *myo-3* promoter. In many models of neurotoxic proteins, expression in muscle cells causes larval arrest and/or motility defects [32, 38]. We found that muscle expression of (Leu)_38_-GFP caused a highly penetrant larval arrest phenotype, whereas expression of (Q)_38_-GFP, (Gln)_38_-GFP, (Cys)_37_-GFP, and (Ala)_38_-GFP caused a weakly penetrant larval arrest phenotype (Figure 2A). (Ser)_38_-GFP did not induce larval arrest. We also examined the effect of each RAN polypeptide on age-dependent post-developmental muscle function, using a previously described conditional expression approach [20]. (Q)_38_-GFP caused a significant increase in paralysis of animals as they aged. However, (Gln)_38_-GFP, (Leu)_38_-GFP, and (Ala)_38_-GFP caused no significant enhancement in paralysis (Figure 2B). To more quantitatively measure changes in motility, we performed thrashing assays. While only (Q)_38_-GFP caused age-dependent paralysis, (Q)_38_-GFP, (Cys)_37_-GFP, (Leu)_38_-GFP, and (Ala)_38_-GFP all caused a decrease in thrashing rates (Figure 2C). (Ala)_38_-GFP motility defects were highly penetrant, whereas (Q)_38_-GFP, (Cys)_37_-GFP, and (Leu)_38_-GFP defects were more variable. Similar to motor neuron expression, muscle expression of (Q)_38_-GFP, but not of codon-varied (Gln)_38_-GFP, caused a thrashing defect (Figure 2C). To test if repeat length affected toxicity, we increased the number of repeats from 38 to 90. Although we were unable to synthesize 90 repeats of polyQ or codon-varied polyAla for unknown technical reasons, (Gln)_90_-GFP and (Leu)_90_-GFP caused significant thrashing defects (Figure 2D) while (Ser)_90_-GFP and (Cys)_90_-GFP did not. None of the HD polypeptides caused a shortening of lifespan when expressed at 38 repeats (Figure 2E). Together, these data in muscle suggest that factors other than polyglutamine contribute to the toxicity of polyQ.

**Figure 2.**
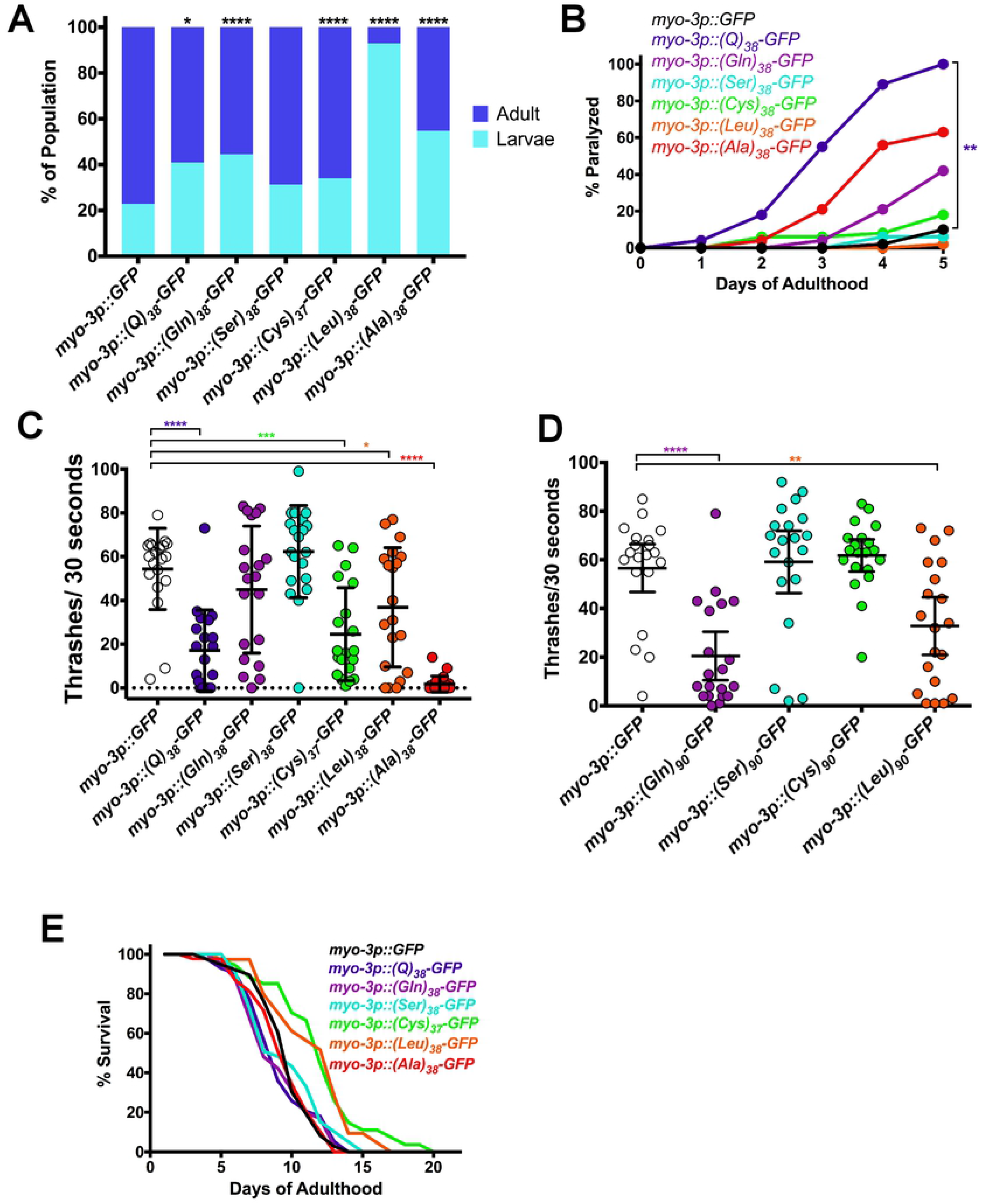
PolyLeu is toxic in muscle. (A) Larval arrest measured by COPAS Biosorter. N = 188-269 animals/genotype. *P<0.05, ****P<0.0001 versus GFP control (Kruskal Wallis test of multiple comparisons with Dunn’s post hoc multiple comparison test). (B) Paralysis assay for adult animals raised in the absence of *gfp*(RNAi). N = 50 animals/genotype. **P<0.01 versus GFP control (Log Rank Test with Bonferroni-adjusted p-value). (C) Liquid thrashing quantification of transgenic animals raised in the absence of *gfp*(RNAi), expressing the indicating polypeptides under the muscle-specific *myo-3* promoter. N = 20 animals/genotype. Each symbol represents one animal, the horizontal line is the mean and the bars represent the standard deviation. *P<0.05, ***P<0.001, ****P<0.0001 versus GFP control (one-way non-parametric ANOVA with Dunn’s post hoc test). (D) Liquid thrashing quantification of transgenic animals expressing the indicated polypeptides under the muscle-specific *myo-3* promoter. N = 20 animals/genotype. **P<0.01, ****P<0.0001 versus GFP control (one-way non-parametric ANOVA with Dunn’s post hoc test). (E) Lifespan measured in transgenic animals raised in the absence of *gfp*(RNAi), expressing the indicating polypeptides under the muscle-specific *myo-3* promoter. N = 50 animals/genotype. P>0.05 for all versus GFP control. (Log Rank Test with Bonferroni-adjusted P-value).

### Localization Patterns of HD RAN Polypeptides

To gain insights into the cell biological properties of the HD RAN polypeptides, we took advantage of the transparent nature of *C. elegans* and performed live animal fluorescent imaging. Previous studies found that polyQ and polySer form puncta at disease-relevant repeat lengths [22, 38]. However, the localization properties of the other codon-varied RAN polypeptides have not been reported. Both (Q)_38_-GFP and (Gln)_38_-GFP had both diffuse signal and puncta, consistent with previous polyQ models (Figure 3A). (Ser)_38_-GFP formed puncta with no detectable diffuse signal (Figure 3A), consistent with previous polySer reports [22]. (Ala)_38_-GFP and (Cys)_37_-GFP also formed puncta. (Cys)_90_-RFP and (Ser)_90_-GFP exhibited strong co-localization when co-expressed, although they did not co-localize with a previous polyQ protein model [24], suggesting that these peptides exist in a structural state that does not allow interactions with polyQ (Figure 3B). Unlike the other RAN polypeptides, the GFP signal for (Leu)_38_-GFP was only detectable in vulval muscle cells and was not detectable in body wall muscle cells (Figure 3A). The low polyLeu signal could be due to localization within a cellular compartment that impairs GFP fluorescence. GFP has impaired folding in oxidizing environments such as the lumen of the endoplasmic reticulum (ER) [39]. Additionally, leucine repeats commonly insert into membranes [40, 41]. Therefore, one possibility is that (Leu)_38_-GFP is in a membrane and the GFP tag is oriented within an oxidizing environment. To test this possibility, we replaced GFP with superfolder GFP (sfGFP), which has several point mutations that enhance folding and fluorescence in non-optimal environments, such as the ER [42, 43]. Unlike (Leu)_38_-GFP, (Leu)_38_-sfGFP was observed in both adult muscle cells and vulval muscle cells and localized to the periphery of large spherical bodies of unknown origin (Figure 3C). The ability to visualize (Leu)_38_-sfGFP but not (Leu)_38_-GFP is consistent with the possibility that (Leu)_38_ may be membrane-bound and localized to an environment that impairs folding of GFP.

**Figure 3.**
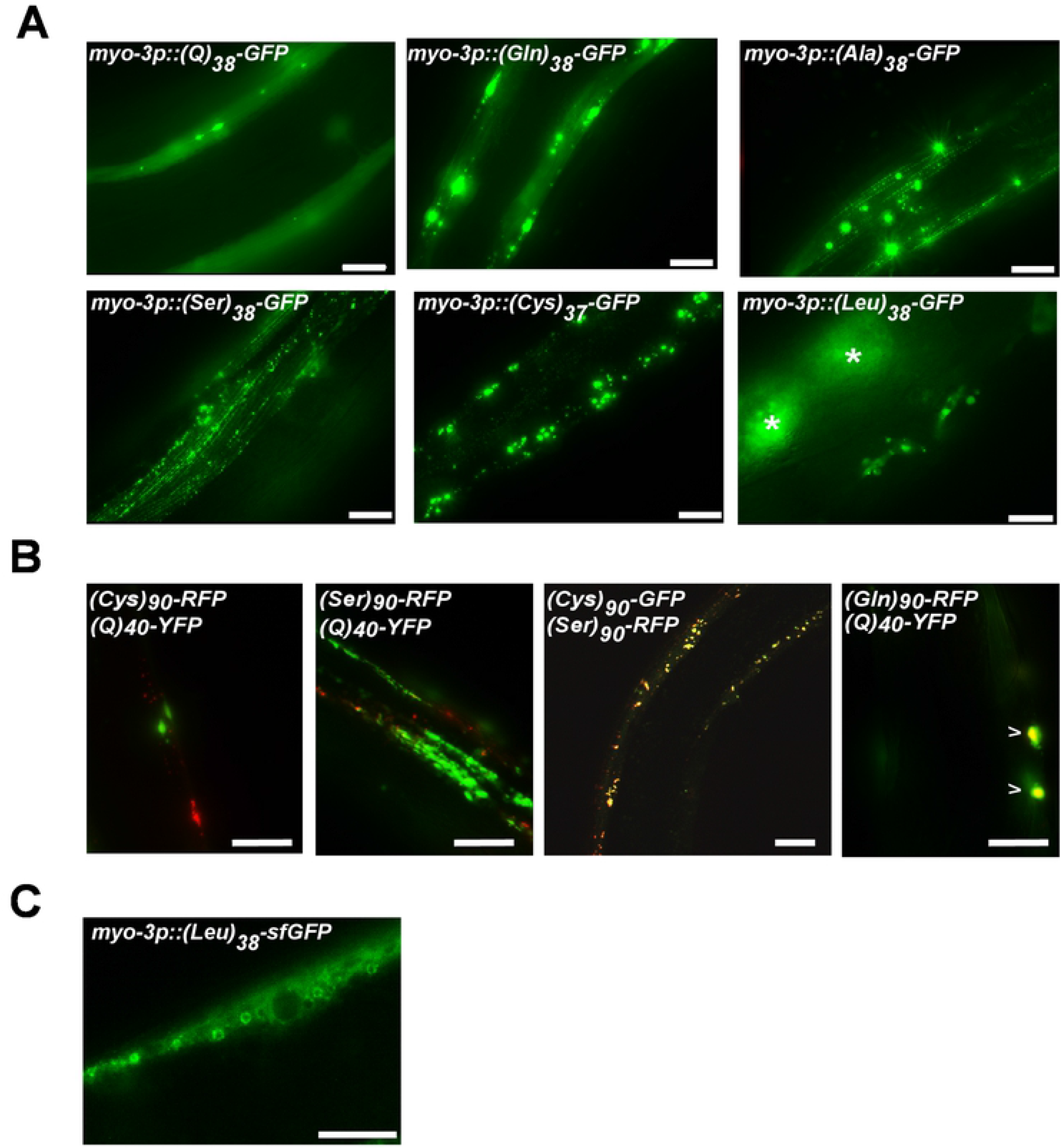
Non-polyGln CAG-associated RAN polypeptides form puncta at ≥38 repeats that are spatially distinct from polyQ aggregates. (A) Representative images of the indicated codon-varied polypeptide in muscle expressed under *myo-3* promoter. Asterisks mark intestinal autofluorescence. Images are from day 1 adults. Scale bar = 10 μm. (B) Co-expression of the indicated RAN polypeptides at 90 repeats with either other RAN polypeptides or (Q)_40_-YFP expressed in AM141. Scale bar = 10 μm. (C) Representative image of *myo-3p::(Leu)_38_-sfGFP*. Scale bar = 10 μm.

### All HD RAN Polypeptides Form Aggregates

PolyGln and polySer are known to form bona fide protein aggregates [22, 44]. However, the structure of the puncta containing the other HD RAN products is unknown. We used Fluorescence Recovery After Photobleaching (FRAP), to better characterize the HD RAN puncta [45]. We found that polyGln, polySer, polyCys, and polyAla puncta each exhibited limited FRAP recovery similar to previously characterized aggregated polyQ (Figure 4). PolyLeu FRAP recovery was more rapid and extensive, suggesting that polyLeu structures have more freely diffusible molecules within the puncta and/or more exchange of molecules with the surrounding cytosol. The total recovery of (Leu)_38_ was ~20%, compared to <10% recovery for the other RAN polypeptides, suggesting that polyLeu may be toxic by affecting different cellular pathways than polyglutamine or other HD RAN polypeptides.

**Figure 4.**
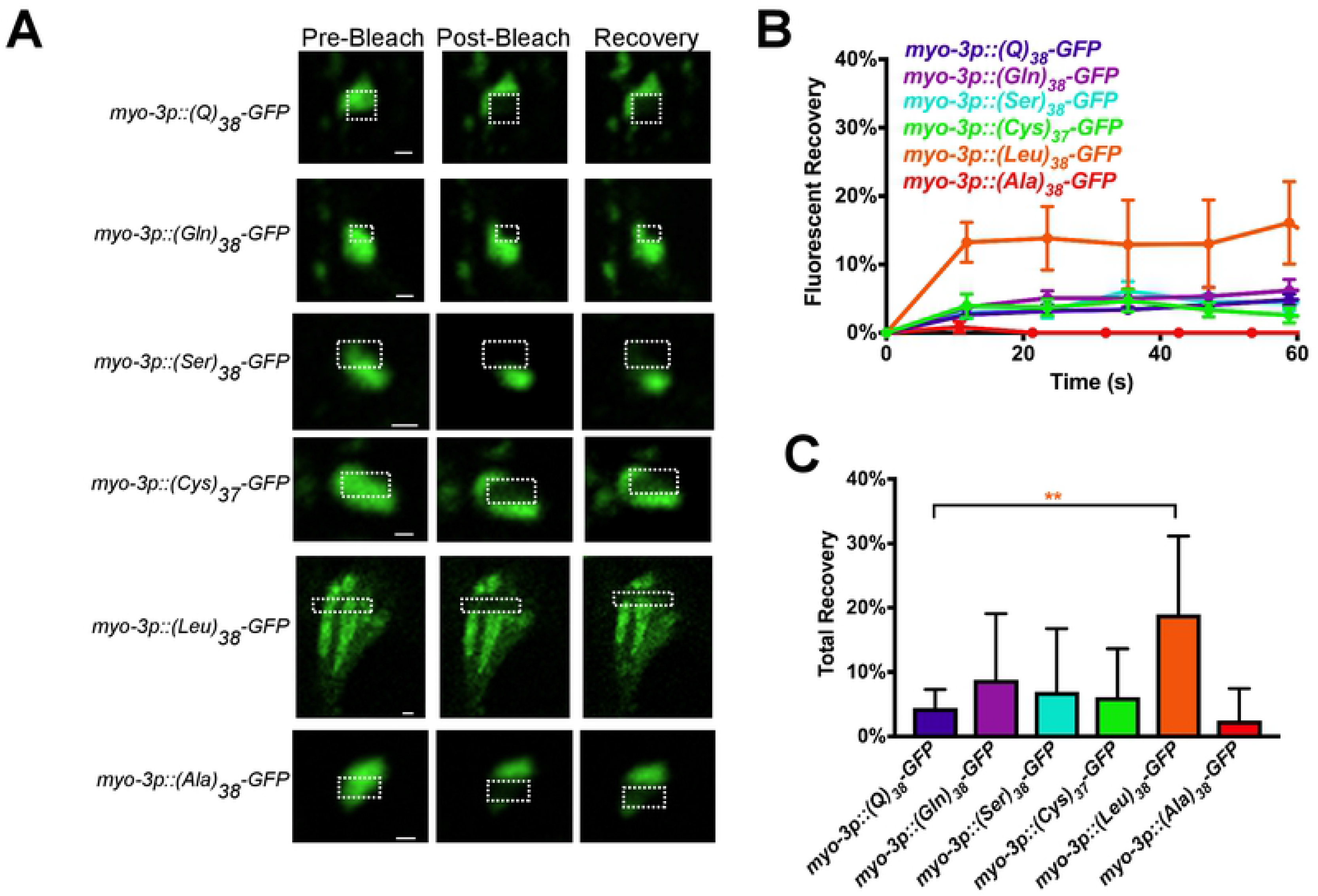
PolyLeu forms protein aggregates with biophysical properties that differ from other CAG-derived RAN polypeptides. (A) Representative images from FRAP analysis of codon-varied RAN polypeptides expressed in muscle. Dashed outline indicates the site of photobleaching and postbleaching quantification. Recovery images are 60 seconds post-bleach. Scale bar =1 μm. (B) Quantification of FRAP imaging. Data shown are mean ± SEM. N = 5-17 puncta/genotype. (C) Average equilibrium fluorescence after 60 seconds of recovery. Data shown are mean ± SEM for 5-17 puncta. **P<0.01 versus polyQ control (one-way non-parametric ANOVA with Dunn’s post hoc test).

### PolyLeu Toxicity is Length Dependent

Length-dependent toxicity is a classic characteristic of CAG repeats. PolyGln is well established to exhibit length-dependent toxicity, which we confirmed using our codon-varied constructs (Figure 2C,D). To determine if polyLeu also exhibits length-dependent toxicity, we expressed polyLeu at 11, 20, 29, and 38 repeats in GABAergic neurons and muscle cells. PolyLeu repeats of 29 or below caused no functional defects in GABAergic neurons, as measured by directional reversals (Figure 5A). However, (Leu)_38_-GFP caused a significant decrease in reversal ability (Figure 5A). Therefore, polyLeu exhibits length-dependent toxicity in GABAergic neurons and requires >29 repeats to cause toxicity. The change in toxicity could be due to altered localization patterns of polyLeu depending on the length of leucine repeats. To test this hypothesis, we expressed the various lengths of polyLeu in muscle cells and visualized their localization. (Leu)_11_-GFP localized to reticular structures that filled the muscle cell, as well as small spherical structures ~1 μm in diameter (Figure 5D). (Leu)_20_-GFP localized to the periphery of the muscle cell, potentially the plasma membrane, as well as various structures inside the cell that failed to exhibit a consistent morphology (Figure 5D). Cellular localization of (Leu)_29_-GFP and (Leu)_38_-GFP was challenging to determine since both lengths of polyLeu appeared to inhibit the fluorescence of the GFP attached to the repeats (Figure 5B,D). The length-dependent decrease in fluorescence was specific to the polyLeu-bound GFP, as free RFP expressed in the same tissues did not exhibit a length-dependent decrease in fluorescence (Figure 5C). (Leu)_11_-GFP, (Leu)_20_-GFP, and (Leu)_29_-GFP did not exhibit the larval arrest observed in both (Leu)_38_-GFP and (Leu)_38_-sfGFP. Together, these data suggest that the strong toxicity of (Leu)_38_ is length dependent, since more than 29 leucine repeats are required to cause larval arrest when expressed in muscle, or reversal defects when expressed in motor neurons. The length-dependent and tissue-independent toxicity of polyLeu suggests that polyLeu could contribute to CAG repeat toxicity, either alone or in combination with polyGln.

**Figure 5.**
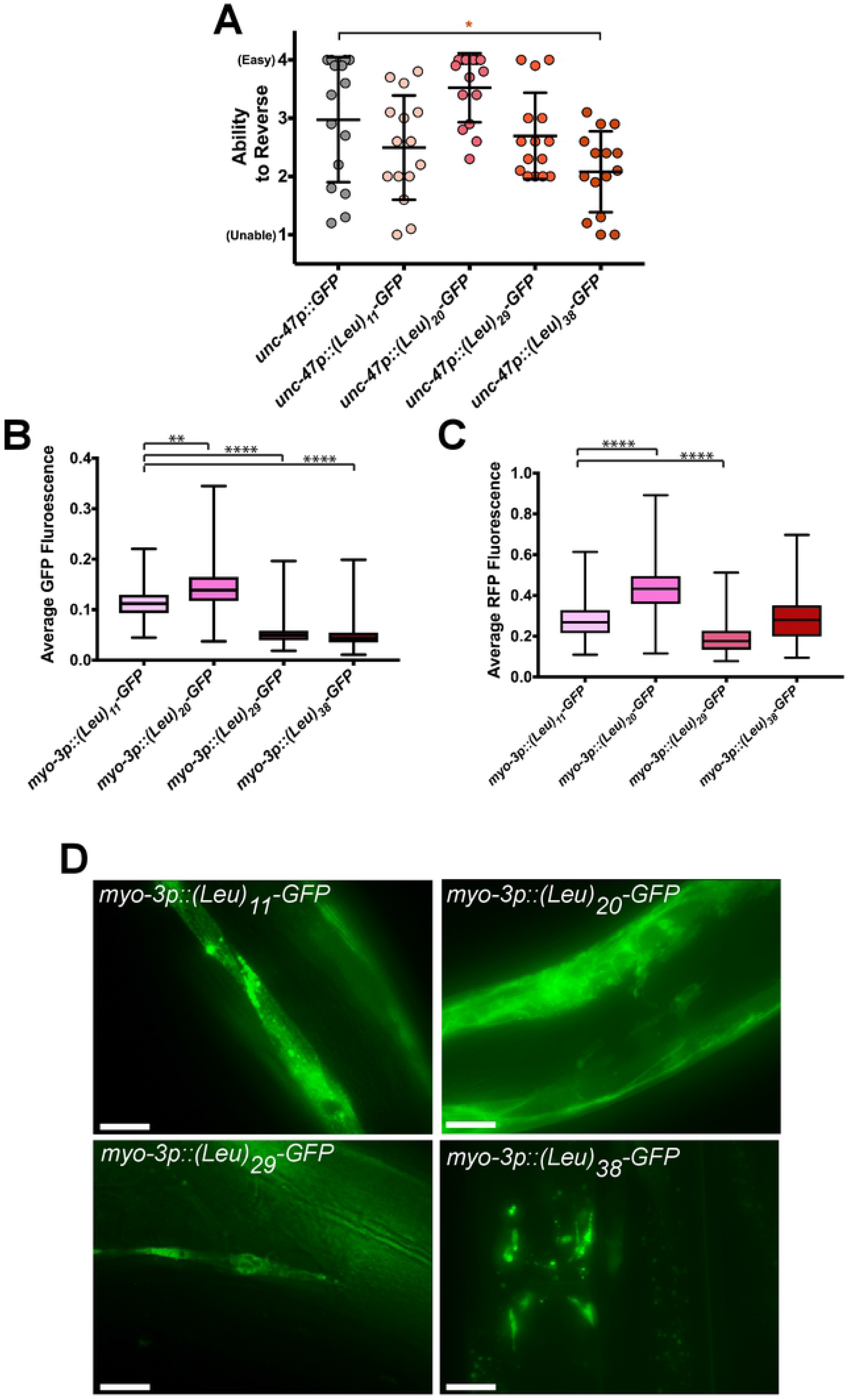
PolyLeu toxicity is length dependent. (A) Quantification of the reversal ability of transgenic animals expressing the indicated polyLeu length in GABAergic neurons. N = 15 animals/genotype. Each symbol represents one animal, the horizontal line is the mean and the bars represent the standard deviation. *P<0.05 versus GFP control (one-way non-parametric ANOVA with Dunn’s post hoc test). (B) Average normalized fluorescence of GFP and (C) RFP in single animals as measured by the COPAS BioSorter. The total GFP or RFP fluorescence was normalized to worm size (time of flight, TOF). N = 149-611 animals/genotype. **P<0.01, ***P<0.001, ****P< 0.0001 versus *myo-3p*::(Leu)_11_-GFP control (one-way non-parametric ANOVA with Dunn’s post hoc test). (D) Representative images of the indicated length of muscle-expressed polyLeu in day 1 adults. Scale bars = 20 μm.

## Discussion

In this study, we investigated the cell biological and pathophysiological properties of the newly discovered HD RAN polypeptides. We found that all of the HD RAN polypeptides formed biophysically defined protein aggregates. However, only a single HD RAN polypeptide, polyLeu, caused both neuropathological changes and functional defects. Given that these studies represent the first *in vivo* dissection of CAG-associated RAN polypeptide properties, we discuss how our findings integrate with previous reports of the structure and function of each homopolymeric peptide.

### CAG Repeats Exhibit Polyglutamine-Independent Toxicity

The pure CAG-encoded polyglutamine (polyQ) and the codon-varied polyglutamine (polyGln) exhibit different levels of toxicity in our model. GABAergic expression of (Q)_38_ caused significant structural and functional neuronal defects, but GABAergic expression of codon-varied (Gln)_38_ caused no detectable defects. Similarly, muscle expression of (Q)_38_ caused extensive motility defects, while muscle expression of codon-varied (Gln)_38_ caused no motility defects. PolyGln did display length-dependent toxicity, as muscle expression of (Gln)_90_ produced motility defects. However, shorter, disease-relevant lengths of polyGln did not cause toxicity, while shorter, disease-relevant lengths of polyQ did cause toxicity.

The disparity in the toxicity of these two ‘polyglutamine’ models is antithetical to the widely-held belief that polyglutamine is the driving cause of the toxicity in HD. Our data suggest that factors other than the translation of CAG repeats into a polyglutamine protein can underlie CAG repeat toxicity. While at odds with dogma, these findings are consistent with recent human genetic studies suggesting that CAG repeat length, but not polyglutamine repeat length, correlates with the age of disease onset in HD [11, 12]. Such CAG-dependent, polyglutamine-independent factors could include RNA repeat toxicity [46] or RAN polypeptide toxicity [22]. CAG repeat models in many organisms, including *C. elegans*, are synonymously referred to as ‘polyQ’ or ‘polyglutamine’ models. However, at least some of these models undergo RAN translation and produce RAN polypeptides in addition to polyglutamine [10]. In the future, it will be important to re-examine whether or not existing *C. elegans* CAG models also undergo RAN translation and produce RAN polypeptides in addition to polyGln. The presence of such polypeptides may dramatically alter the use and interpretation of studies based on these models. Given the widespread prevalence of RAN translation in several CAG repeat expansion disorders, referring to these models solely as ‘polyQ models’ may no longer be accurate and our descriptions should be modified to include our evolving understanding of the role of RAN polypeptides in addition to polyQ.

### PolyCys and PolySer: Weakly Toxic RAN Products

Both polyCys and polySer exhibited similar cell biological and phenotypic effects when expressed in *C. elegans*. PolyCys and polySer formed aggregates at 38 and 90 repeats. They also co-localized when expressed together. The colocalization of polySer and polyCys is not solely due to their polar charge. Glutamine is also a polar amino acid, but polyGln did not colocalize with either polySer or polyCys (Figure 3B). A possible explanation of the distinct aggregation patterns of polyGln, polySer, and polyCys could be that polySer and polyCys may localize to a different subcellular environment than polyGln. For example, both Cys and Ser [47], but not Gln, are known targets for post-translational protein palmitoylation, which anchors proteins to cellular membranes [48]. The palmitoylation of Cys residues in another protein, cysteine string protein α, is essential for protein aggregation and is associated with the neurodegenerative disease adult-onset neuronal ceroid lipofuscinosis [49]. Therefore, post-translational modification of polySer and polyCys, but not polyGln, may lead to protein interactions that drive protein aggregation in ways that are distinct from polyGln. Future studies examining the post-translational modification state of the polySer and polyCys protein could provide insights into these and other potential aggregation mechanisms.

Another common property of polyCys and polySer is that expression of either polypeptide caused weak and inconsistently toxic effects. GABAergic expression of (Cys)_37_ caused minor structural but not functional defects, whereas GABAergic expression of (Ser)_38_ caused functional defects, but not structural defects. Muscle expression of (Cys)_37_ caused weak motility defects and larval arrest. However, muscle expression of polySer never caused motility defects or larval arrest. When expressed on their own, polyCys and polySer exhibit little to no phenotypic consequences in *C. elegans*. This was surprising considering a recent study which suggested that polySer has significant toxicity in a SCA8 murine model, another CAG repeat expansion disease [50]. However, this model did not express a pure polySer protein. Rather, it expressed the CAG expanded ATXN8 gene which is the genetic cause of SCA8. PolySer is one of several RAN products translated from the SCA8 CAG repeat. However, the other possible RAN products were not measured. In the murine model of SCA8, polySer was thought to be toxic because RAN translated polySer colocalized with degenerating brain regions which lacked the ATXN8 protein. However, RAN translation products from different reading frames typically occur in the same tissue [22], and can occur in the same cells [51]. Therefore, polySer may be acting as a biomarker for RAN translation, and one or more of the other CAG RAN products are toxic in neurons. Our studies suggest that polySer alone does not cause neurodegeneration. However, polySer may have synergistic interactions with other RAN polypeptides that enhance toxicity. In the future, it will be important to test this possibility by coexpressing polySer with each of the other RAN products to determine if there are synergistic effects on toxicity. In addition, it will also be important to place polySer, as well as other RAN polypeptides, in a more endogenous genetic context by including downstream sequence appropriate to each RAN polypeptide reading frame.

### PolyAla: A Possible Contributor to CAG Toxicity

PolyAla may be a strong contributor to CAG repeat toxicity. In *C. elegans*, polyAla was toxic in both muscle and GABAergic neurons. Muscle expression of polyAla caused strong motility defects and GABAergic expression of polyAla caused functional, but not structural, defects. (Ala)_38_-GFP also formed aggregates when expressed in muscle cells of *C. elegans*. In humans, expanded polyAla repeats cause aggregation in multiple diseases [52]. The aggregation of the polyAla repeats caused the mis-localization of the polyAla containing proteins and a concordant loss-of-function phenotype [52]. Since polyAla toxicity is mediated by the loss-of-function of the protein containing the expanded Ala tract, polyAla toxicity has mainly been studied within its native genetic context. The toxicity of a polyAla peptide without genetic context, as was modeled in our study, has not been thoroughly studied. In addition, because the expanded alanine repeats in the polyAla diseases are on average shorter than 30 repeats, a pure polyAla peptide had not been modeled at an HD relevant length until our work.

A homopolymeric alanine peptide may be toxic through the similar mechanisms as the alanine repeat-expansion diseases. A yeast-two hybrid model found that homopolymeric disease-relevant lengths of polyAla ((Ala)_29_ and (Ala)_28_) exhibit self-interactions [53]. It is unknown if alanine repeat lengths, which normally appear in proteins (5-21 repeats) [54], also interact with disease-relevant lengths of alanine repeats. PolyAla repeats are enriched in transcription factors [54, 55], which need to be localized to the nucleus to perform their function. Therefore, interaction of the transcription factors containing polyAla repeats with the polyAla aggregates could deplete the cell of available transcription factors. PolyAla aggregates sequestering transcription factors could also explain the difference in toxicity observed between muscle cells and GABAergic neurons, as the depleted transcription factors may be less important in GABAergic neurons than in muscle cells.

### PolyLeu Is the Most Toxic HD RAN Product

In our *C. elegans* CAG RAN polypeptide models, the most toxic HD RAN product across tissues was polyLeu. Like polyAla, polySer, and polyCys, polyLeu caused functional defects when expressed in GABAergic neurons and motility defects when expressed in muscle. However, polyLeu expression caused highly penetrant phenotypes in both muscle and neurons which were only weakly observed in the other RAN models. GABAergic expression of polyLeu induced significant morphological defects where every commissure exhibited structural disorganization. Muscle expression of (Leu)_38_-GFP caused larval arrest of ~90% of the population.

The strong toxicity of polyLeu was unexpected, as polyLeu repeat expansions have not been previously linked to a genetic disease. Interestingly, polyLeu repeats are one of the most common single amino acid repeats in the human proteome, with ~1,500 proteins containing polyLeu repeats at least four leucines [54]. However, none of the naturally occurring polyLeu repeats exceed 11 leucine repeats, suggesting there is an evolutionary selection against longer polyLeu repeats [54]. The lack of polyLeu repeat expansion diseases could be due to polyLeu repeat expansions disrupting development in humans, as it does in *C. elegans*. In mammalian cells, RAN translation rates, which produce polyLeu, increase with activation of the integrated stress response pathway [14, 16, 56], and the integrated stress response pathway is activated with age [57], suggesting that RAN translation rates could increase with age. If RAN translation is correlated with age, RAN-translated polyLeu would not affect development. Another possible reason for the lack of diseases caused by polyLeu repeat expansions is that polyLeu is produced from six different codons, while glutamine is encoded by only two codons. Therefore, codon heterogeneity may effectively protect against repetitive polyLeu-encoding nucleotide sequences. Because polyLeu repeat sequences are not commonly associated with disease, they have received little attention. Limited previous work found that polyLeu repeats longer than 30 repeats caused toxicity in cells [58, 59] and in *Drosophila* [60], but these models were created with a CTG repeat that could be a substrate for RAN translation and subsequent production of other peptides. Our work is the first evidence that a codon-varied pure polyLeu peptide causes significant cellular toxicity.

The toxicity of polyLeu could be due to the apparent membrane-localization of polyLeu. While the expression pattern of polyLeu changes based on repeat length, all lengths appear to be membrane localized. This is consistent with previous findings that polyLeu peptides as short as nine repeats can spontaneously incorporate into lipid membranes [40, 61, 62]. Leucine occurs in many signal peptides and transmembrane domains [55]. For this reason, leucine is the most common amino acid in proteins that localize to the Endoplasmic Reticulum (ER), Golgi apparatus, or vacuoles [63]. Consistent with this, (Leu)_11_-GFP localizes to a tubular network in *C. elegans* that resembles previous reports of ER [64] and spherical bodies ~1.2 μm in diameter which cluster at the distal regions of the muscle cells. Eleven leucine repeats can act as an anchor signal sequence to target proteins to a membrane [62], so the spherical structures may be part of the endocytic pathway. (Leu)_20_-GFP appears to localize to the plasma membrane and smaller membrane-bound compartments based on its localization to structures similar to the peripheral shape of a *C. elegans* muscle cell, as well as other spherical and non-spherical structures. Although we were unable to detect (Leu)_38_-GFP, (Leu)_38_-sfGFP localized to the boundaries of unidentified spherical structures. These structures are distributed throughout the length of the muscle cell and are similar in size to those seen with (Leu)_11_. (Leu)_38_-sfGFP also has a diffuse background signal which may represent ER localization, since this signal was visible with sfGFP-tagged (Leu)_38_ but not GFP-tagged (Leu)_38_. Whether or not (Leu)_38_ disrupts ER homeostasis and how this may contribute to polyLeu toxicity will be the focus of future work.

In conclusion, we have characterized the *in vivo* properties of novel CAG-associated RAN homopolymers in *C. elegans*. Our findings suggest that CAG-derived polypeptides other than polyglutamine can cause toxicity through mechanisms that are likely independent of polyQ. While much remains to be learned about these new polypeptides, our findings and those of others strongly suggests that CAG repeat disease therapies targeting only polyglutamine toxicity mechanisms may not be effective. Instead, approaches that mitigate the toxicity of other CAG RAN polypeptides, particularly polyleucine, may need to be considered.

## Materials and methods

### *Caenorhabditis elegans* strains and culture

Strains were cultured on standard NGM media with *gfp*(RNAi) bacteria at 20°C until the generation before the experiment. For motility assays, animals were picked at the L4 stage, shifted to *E. coli* OP50, and allowed to have progeny at 20°C. The progeny were picked as L4 animals, kept on *E. coli* OP50, and placed at 25°C. All experiments were performed at 25°C.

### Molecular biology and transgenics

Codon-varied homopolymer sequences were synthesized for the 90 repeat polypeptides (Integrated DNA Technologies, Coralville, Iowa, USA). 38 repeat polypeptides were made using a “building block” approach where codon-varied sequences for 11 repeats were synthesized (GeneWiz, South Plainfield, NJ, USA). Each “building block” could be extended by digesting the vector containing a building block with BsmbI, and digesting the insert containing a building block with BsaI as previously described [65]. The nucleotide sequences used for the HD polypeptides are listed in Table S1. Promoters were produced and cloned in as previously described [93].

Transgenic worms were generated by injecting the RAN polypeptide construct (20 ng/μl) and the *myo-3p::mCherry pCFJ104* marker plasmid (100 ng/μl) into the gonad of wild-type animals. Transgenes were integrated using a standard gamma ray (Cs^137^) mutagenesis, followed by selection of animals exhibiting 100% transmission of the mCherry marker. Integrated strains were outcrossed six times to wild-type animals. Injected animals were maintained on *gfp*(RNAi) plates until the experimental assay was performed. All procedures involving recombinant or synthetic nucleic acid molecules and materials were approved by the University of Pittsburgh Institutional Biosafety Committee.

### Reversal assays

30-40 progeny were picked as L4 animals and moved to 25°C on *E. coli* OP50 bacteria. 24 hours later, the progeny were tested for their ability to reverse (N=20 animals/genotype). All assays were performed with the experimenter blinded to genotype. Animals were lightly tapped on the head with a platinum pick and scored from 1-4 on their ability to reverse as previously described [66]. Each worm was scored 5 consecutive times, as wild-type worms began to acclimate after 7 taps.

### Commissure assays

Strains were moved from 20°C to 25°C at least a generation before the experiment and maintained on *E. coli* OP50. L4 animals of the indicated genotype were isolated at 25°C and imaged 24 hours later as ‘Day 1 adults’. All of the strains contained an *unc-47p::RFP* marker to reveal GABAergic motor neuron morphology. Animals were anesthetized in 10 mM levamisole and Z-series images of GABAergic commissures were collected. Commissure breaks were identified as interruptions in the RFP signal surrounded by dorsal and ventral RFP in the commissures as previously described [20]. Blebbing was scored only in the commissures and was identified by the presence of one or more RFP varicosities as previously described [20]. Abnormal commissures were identified as those exhibiting branching and/or failing to reach the dorsal side of the animal.

### Microscopy

Day 1 adult *C. elegans* were anesthetized in 10 mM levamisole for 10 minutes. The animals were then mounted on 3% agarose pads for fluorescence microscopy. Wide field images were collected on a Leica DMi4000 inverted microscope and a Leica DFC 340x digital camera (Leica Microsystems, Wetzlar, Germany). Z-stack images were deconvolved using Leica AF6000 software.

For FRAP studies, confocal fluorescence images were captured on a Nikon A1plus confocal microscope through an Apo 60x/ 1.4NA Oil objective lens. The microscope was operated on the NIS-Elements AR version 5.02 software platform. GFP or RFP were excited at 488 or 561, respectively. Images were captured every 11 seconds to avoid bleaching over the course of imaging. Following imaging of baseline fluorescence, a region of interest corresponding to a portion of the puncta was photobleached and fluorescence recovery within the photobleached area was monitored over at least 60 seconds. Data were normalized so that the image preceding the photobleach was set to 100% and the first image following the photobleach was set to zero percent. Imaging conditions over the time course of the experiment caused minimal loss of signal, suggesting an absence of photobleaching during the monitoring period. FRAP analysis was performed using Fiji software [67]. X-Y drift was corrected using the “Correct 3d Drift” plugin [68].

### COPAS experiments

Gravid animals grown on *gfp*(RNAi) were placed on *E. coli* OP50 for 6 hours at 20°C to collect a synchronized brood. The adults were then removed and the plates were placed at 25°C. 48 hours later, the progeny were sorted through a COPAS BioSorter (Union Biometrica, Holliston, MA, USA). Worm time of flight (TOF) was measured. Animals with a TOF ≥ 300 were counted as adults. Animals with a TOF < 300 were counted as larvae. When measuring GFP and RFP, fluorescent detection settings were identical for all samples. Average fluorescence was measured as total fluorescence divided by the time of flight.

### Paralysis assays

Gravid animals were moved from *gfp*(RNAi) to *E. coli* OP50 and allowed to lay eggs for 24 hours at 20°C. The resulting progeny were allowed to grow up on *E. coli* OP50, permitting RAN polypeptide accumulation. Ten L4 animals were placed on each of the five-3cm plates spotted with *E. coli* OP50 (N=50 animals/genotype) and moved to 25°C. Each day, animals that failed to move at least half a body length in response to manual stimulation with a platinum wire but were still alive (pharyngeal pumping, movement of less than half a body length) were scored as paralyzed. Animals that died, desiccated on the plate edges, or exhibited internal hatching of progeny were censored from the assay. Each day, mobile animals were transferred to a new plate and paralyzed, dead, or censored animals were removed from the assay.

### Thrashing assays

Gravid animals were moved from *gfp*(RNAi) to *E. coli* OP50 and allowed to lay eggs for 24 hours at 20°C. The resulting progeny were allowed to grow up on *E. coli* OP50, permitting polypeptide accumulation. The day before the experiment, 40 transgenic L4 animals were transferred to an *E. coli* OP50 plate and placed at 25°C. The following day, worms were placed on clean NGM plates and allowed to move freely for 10 minutes to remove excess bacteria. Worms were then placed individually into 3cm petri dishes containing M9 buffer and allowed to adjust to the new environment for 5 minutes. The worms were observed for 30 seconds and the number of thrashes (reversal of body bend that crosses the midline) was counted.

### Lifespan assays

Gravid animals were moved from *gfp*(RNAi) to *E. coli* OP50 and allowed to lay eggs for 24 hours at 20°C. The resulting progeny were allowed to grow up on *E. coli* OP50, permitting polypeptide accumulation. 10 L4 animals were transferred to five-3cm plates (N=50 animals/genotype). Lifespan assays were performed at 25°C with *E. coli* OP50 spotted on NGM plates. Worms were classified as alive, dead (no movement in response to touch with a wire), or censored (lost or bagged worms) once per day for lifespan assays.

### Statistical analysis

Comparison of means were analyzed with ANOVA using Dunn’s post hoc test analysis in GraphPad Prism 7 (GraphPad Software, Inc., La Jolla, CA, USA). For the FRAP analysis, the plateau was measured using one phase association in GraphPad Prism 7. Paralysis assay and lifespan assays were analyzed using the Kaplan-Meier logrank function (OASIS) [69]. P-values of <0.05 were considered to be significant.

## Acknowledgements

This work was supported by grants from the NIH (R21NS112617, T.L.). Some strains were provided by the CGC, which is funded by NIH Office of Research Infrastructure Programs (P40 OD010440).

## Conflict of Interest Statement

The authors have no conflicts of interests to declare

## References

1. Paulson H. Repeat expansion diseases. Handbook of clinical neurology. 2018;147:105–23. Epub 2018/01/13. doi: 10.1016/B978-0-444-63233-3.00009-9. PubMed PMID: 29325606; PubMed Central PMCID: PMCPMC6485936.

2. Stenoien DL, Cummings CJ, Adams HP, Mancini MG, Patel K, DeMartino GN, et al. Polyglutamine-expanded androgen receptors form aggregates that sequester heat shock proteins, proteasome components and SRC-1, and are suppressed by the HDJ-2 chaperone. Human molecular genetics. 1999;8(5):731–41. PubMed PMID: 10196362.

3. Lee WC, Yoshihara M, Littleton JT. Cytoplasmic aggregates trap polyglutamine-containing proteins and block axonal transport in a Drosophila model of Huntington’s disease. Proceedings of the National Academy of Sciences of the United States of America. 2004;101(9):3224–9. doi: 10.1073/pnas.0400243101. PubMed PMID: 14978262; PubMed Central PMCID: PMC365771.

4. Park SH, Kukushkin Y, Gupta R, Chen T, Konagai A, Hipp MS, et al. PolyQ proteins interfere with nuclear degradation of cytosolic proteins by sequestering the Sis1p chaperone. Cell. 2013;154(1):134–45. Epub 2013/06/25. doi: 10.1016/j.cell.2013.06.003. PubMed PMID: 23791384.

5. Alves-Rodrigues A, Gregori L, Figueiredo-Pereira ME. Ubiquitin, cellular inclusions and their role in neurodegeneration. Trends in neurosciences. 1998;21(12):516–20. Epub 1999/01/09. PubMed PMID: 9881849.

6. Saudou F, Finkbeiner S, Devys D, Greenberg ME. Huntingtin acts in the nucleus to induce apoptosis but death does not correlate with the formation of intranuclear inclusions. Cell. 1998;95(1):55–66. Epub 1998/10/20. doi: 10.1016/s0092-8674(00)81782-1. PubMed PMID: 9778247.

7. Ashkenazi A, Bento CF, Ricketts T, Vicinanza M, Siddiqi F, Pavel M, et al. Polyglutamine tracts regulate beclin 1-dependent autophagy. Nature. 2017;545(7652):108–11. doi: 10.1038/nature22078. PubMed PMID: 28445460; PubMed Central PMCID: PMC5420314.

8. Berendzen KM, Durieux J, Shao LW, Tian Y, Kim HE, Wolff S, et al. Neuroendocrine Coordination of Mitochondrial Stress Signaling and Proteostasis. Cell. 2016;166(6):1553–63 e10. doi: 10.1016/j.cell.2016.08.042. PubMed PMID: 27610575.

9. Nollen EA, Garcia SM, van Haaften G, Kim S, Chavez A, Morimoto RI, et al. Genome-wide RNA interference screen identifies previously undescribed regulators of polyglutamine aggregation. Proceedings of the National Academy of Sciences of the United States of America. 2004;101(17):6403–8. doi: 10.1073/pnas.0307697101. PubMed PMID: 15084750; PubMed Central PMCID: PMC404057.

10. Banez-Coronel M, Ayhan F, Tarabochia AD, Zu T, Perez BA, Tusi SK, et al. RAN Translation in Huntington Disease. Neuron. 2015;88(4):667–77. doi: 10.1016/j.neuron.2015.10.038. PubMed PMID: 26590344; PubMed Central PMCID: PMC4684947.

11. Genetic Modifiers of Huntington’s Disease Consortium. Electronic address ghmhe, Genetic Modifiers of Huntington’s Disease C. CAG Repeat Not Polyglutamine Length Determines Timing of Huntington’s Disease Onset. Cell. 2019;178(4):887–900 e14. Epub 2019/08/10. doi: 10.1016/j.cell.2019.06.036. PubMed PMID: 31398342.

12. Wright GEB, Collins JA, Kay C, McDonald C, Dolzhenko E, Xia Q, et al. Length of Uninterrupted CAG, Independent of Polyglutamine Size, Results in Increased Somatic Instability, Hastening Onset of Huntington Disease. American journal of human genetics. 2019;104(6):1116–26. Epub 2019/05/21. doi: 10.1016/j.ajhg.2019.04.007. PubMed PMID: 31104771; PubMed Central PMCID: PMCPMC6556907.

13. Banez-Coronel M, Ranum LPW. Repeat-associated non-AUG (RAN) translation: insights from pathology. Lab Invest. 2019;99(7):929–42. Epub 2019/03/29. doi: 10.1038/s41374-019-0241-x. PubMed PMID: 30918326.

14. Westergard T, McAvoy K, Russell K, Wen X, Pang Y, Morris B, et al. Repeat-associated non-AUG translation in C9orf72-ALS/FTD is driven by neuronal excitation and stress. EMBO Molecular Medicine. 2019;11(2):e9423. doi: 10.15252/emmm.201809423.

15. Tabet R, Schaeffer L, Freyermuth F, Jambeau M, Workman M, Lee CZ, et al. CUG initiation and frameshifting enable production of dipeptide repeat proteins from ALS/FTD C9ORF72 transcripts. Nature communications. 2018;9(1):152. Epub 2018/01/13. doi: 10.1038/s41467-017-02643-5. PubMed PMID: 29323119; PubMed Central PMCID: PMCPMC5764992.

16. Cheng W, Wang S, Mestre AA, Fu C, Makarem A, Xian F, et al. C9ORF72 GGGGCC repeat-associated non-AUG translation is upregulated by stress through eIF2α phosphorylation. Nature Communications. 2018;9(1):51-. doi: 10.1038/s41467-017-02495-z. PubMed PMID: 29302060.

17. Zu T, Liu Y, Banez-Coronel M, Reid T, Pletnikova O, Lewis J, et al. RAN proteins and RNA foci from antisense transcripts in C9ORF72 ALS and frontotemporal dementia. Proceedings of the National Academy of Sciences of the United States of America. 2013;110(51):E4968–77. doi: 10.1073/pnas.1315438110. PubMed PMID: 24248382; PubMed Central PMCID: PMC3870665.

18. Nemes JP, Benzow KA, Moseley ML, Ranum LP, Koob MD. The SCA8 transcript is an antisense RNA to a brain-specific transcript encoding a novel actin-binding protein (KLHL1). Human molecular genetics. 2000;9(10):1543–51. Epub 2000/07/11. doi: 10.1093/hmg/9.10.1543. PubMed PMID: 10888605.

19. Zu T, Gibbens B, Doty NS, Gomes-Pereira M, Huguet A, Stone MD, et al. Non-ATG-initiated translation directed by microsatellite expansions. Proceedings of the National Academy of Sciences of the United States of America. 2011;108(1):260–5. doi: 10.1073/pnas.1013343108. PubMed PMID: 21173221; PubMed Central PMCID: PMC3017129.

20. Rudich P, Snoznik C, Watkins SC, Monaghan J, Pandey UB, Lamitina ST. Nuclear localized C9orf72-associated arginine-containing dipeptides exhibit age-dependent toxicity in C. elegans. Human molecular genetics. 2017;26(24):4916–28. Epub 2017/10/17. doi: 10.1093/hmg/ddx372. PubMed PMID: 29036691; PubMed Central PMCID: PMCPMC5886095.

21. Wang ZF, Ursu A, Childs-Disney JL, Guertler R, Yang WY, Bernat V, et al. The Hairpin Form of r(G4C2)(exp) in c9ALS/FTD Is Repeat-Associated Non-ATG Translated and a Target for Bioactive Small Molecules. Cell Chem Biol. 2019;26(2):179–90 e12. Epub 2018/12/06. doi: 10.1016/j.chembiol.2018.10.018. PubMed PMID: 30503283; PubMed Central PMCID: PMCPMC6386614.

22. Bañez-Coronel M, Ayhan F, Tarabochia AD, Zu T, Perez BA, Tusi SK, et al. RAN Translation in Huntington Disease. Neuron. 2015;88:667–77. doi: 10.1016/j.neuron.2015.10.038. PubMed PMID: 26590344.

23. Lee AL, Ung HM, Sands LP, Kikis EA. A new Caenorhabditis elegans model of human huntingtin 513 aggregation and toxicity in body wall muscles. PloS one. 2017;12(3):e0173644. doi: 10.1371/journal.pone.0173644. PubMed PMID: 28282438; PubMed Central PMCID: PMC5345860.

24. Morley JF, Brignull HR, Weyers JJ, Morimoto RI. The threshold for polyglutamine-expansion protein aggregation and cellular toxicity is dynamic and influenced by aging in Caenorhabditis elegans. Proceedings of the National Academy of Sciences of the United States of America. 2002;99(16):10417–22. doi: 10.1073/pnas.152161099. PubMed PMID: 12122205; PubMed Central PMCID: PMC124929.

25. Satyal SH, Schmidt E, Kitagawa K, Sondheimer N, Lindquist S, Kramer JM, et al. Polyglutamine aggregates alter protein folding homeostasis in Caenorhabditis elegans. Proceedings of the National Academy of Sciences of the United States of America. 2000;97(11):5750–5. doi: 10.1073/pnas.100107297. PubMed PMID: 10811890; PubMed Central PMCID: PMC18505.

26. Faber PW, Voisine C, King DC, Bates EA, Hart AC. Glutamine/proline-rich PQE-1 proteins protect Caenorhabditis elegans neurons from huntingtin polyglutamine neurotoxicity. Proceedings of the National Academy of Sciences of the United States of America. 2002;99(26):17131–6. doi: 10.1073/pnas.262544899. PubMed PMID: 12486229; PubMed Central PMCID: PMC139281.

27. Faber PW, Alter JR, MacDonald ME, Hart AC. Polyglutamine-mediated dysfunction and apoptotic death of a Caenorhabditis elegans sensory neuron. Proceedings of the National Academy of Sciences of the United States of America. 1999;96(1):179–84. PubMed PMID: 9874792; PubMed Central PMCID: PMC15113.

28. Graveland GA, Williams RS, DiFiglia M. Evidence for degenerative and regenerative changes in neostriatal spiny neurons in Huntington&#039;s disease. Science. 1985;227(4688):770. doi: 10.1126/science.3155875.

29. Schuske K, Beg AA, Jorgensen EM. The GABA nervous system in C. elegans. Trends in Neurosciences. 2004;27(7):407–14. doi: https://doi.org/10.1016/j.tins.2004.05.005.

30. Garcia SM, Casanueva MO, Silva MC, Amaral MD, Morimoto RI. Neuronal signaling modulates protein homeostasis in Caenorhabditis elegans post-synaptic muscle cells. Genes & Development. 2007;21:3006–16. doi: 10.1101/gad.1575307. PubMed PMID: 18006691.

31. Earls LR, Hacker ML, Watson JD, Miller DM. Coenzyme Q protects *Caenorhabditis elegans* GABA neurons from calcium-dependent degeneration. Proc Natl Acad Sci U S A. 2010;107(32):14460. doi: 10.1073/pnas.0910630107.

32. Rudich P, Snoznik C, Watkins SC, Monaghan J, Pandey UB, Lamitina ST. Nuclear localized C9orf72-associated arginine-containing dipeptides exhibit age-dependent toxicity in C. elegans. Human Molecular Genetics. 2017. doi: 10.1093/hmg/ddx372.

33. Vaccaro A, Tauffenberger A, Aggad D, Rouleau G, Drapeau P, Parker JA. Mutant TDP-43 and FUS Cause Age-Dependent Paralysis and Neurodegeneration in C. elegans. PLOS ONE. 2012;7(2):e31321. doi: 10.1371/journal.pone.0031321.

34. Kraemer BC, Zhang B, Leverenz JB, Thomas JH, Trojanowski JQ, Schellenberg GD. Neurodegeneration and defective neurotransmission in a Caenorhabditis elegans model of tauopathy. Proc Natl Acad Sci U S A. 2003;100(17):9980. doi: 10.1073/pnas.1533448100.

35. Nix P, Hammarlund M, Hauth L, Lachnit M, Jorgensen EM, Bastiani M. Axon Regeneration Genes Identified by RNAi Screening in *C. elegans*. The Journal of Neuroscience. 2014;34(2):629. doi: 10.1523/JNEUROSCI.3859-13.2014.

36. González-Hunt CP, Leung MCK, Bodhicharla RK, McKeever MG, Arrant AE, Margillo KM, et al. Exposure to Mitochondrial Genotoxins and Dopaminergic Neurodegeneration in Caenorhabditis elegans. PLOS ONE. 2014;9(12):e114459. doi: 10.1371/journal.pone.0114459.

37. Kennedy S, Wang D, Ruvkun G. A conserved siRNA-degrading RNase negatively regulates RNA interference in C. elegans. Nature. 2004;427(6975):645–9. doi: 10.1038/nature02302.

38. Morley JF, Brignull HR, Weyers JJ, Morimoto RI. The threshold for polyglutamine-expansion protein aggregation and cellular toxicity is dynamic and influenced by aging in Caenorhabditis elegans. Proc Natl Acad Sci U S A. 2002;99:10417–22. doi: 10.1073/pnas.152161099. PubMed PMID: 12122205.

39. Jain RK, Joyce PB, Molinete M, Halban PA, Gorr SU. Oligomerization of green fluorescent protein in the secretory pathway of endocrine cells. The Biochemical Journal. 2001;360(Pt 3):645–9. PubMed PMID: 11736655.

40. Kuroiwa T, Sakaguchi M, Mihara K, Omura T. Systematic analysis of stop-transfer sequence for microsomal membrane. The Journal of Biological Chemistry. 1991;266(14):9251–5.

41. Chen H, Kendall DA. Artificial Transmembrane Segments.: Requirements for Stop Transfer and Polypeptide Orientation The Journal of Biological Chemistry. 1995;270(23):14115–22.

42. Pédelacq J-D, Cabantous S, Tran T, Terwilliger TC, Waldo GS. Engineering and characterization of a superfolder green fluorescent protein. Nature Biotechnology. 2005;24:79. doi: 10.1038/nbt1172 https://www.nature.com/articles/nbt1172#supplementary-information.

43. Aronson DE, Costantini LM, Snapp EL. Superfolder GFP is fluorescent in oxidizing environments when targeted via the Sec translocon. Traffic (Copenhagen, Denmark). 2011;12(5):543–8. Epub 2011/02/25. doi: 10.1111/j.1600-0854.2011.01168.x. PubMed PMID: 21255213.

44. Satyal SH, Schmidt E, Kitagawa K, Sondheimer N, Lindquist S, Kramer JM, et al. Polyglutamine aggregates alter protein folding homeostasis in Caenorhabditis elegans. Proc Natl Acad Sci U S A. 2000;97:5750–5. doi: 10.1073/pnas.100107297. PubMed PMID: 10811890.

45. Brignull HR, Morley JF, Garcia SM, Morimoto RI. Modeling Polyglutamine Pathogenesis in C. elegans. Methods in Enzymology. 412: Academic Press; 2006. p. 256–82.

46. Hsu R-J, Hsiao K-M, Lin M-J, Li C-Y, Wang L-C, Chen L-K, et al. Long Tract of Untranslated CAG Repeats Is Deleterious in Transgenic Mice. PLOS ONE. 2011;6(1):e16417. doi: 10.1371/journal.pone.0016417.

47. Takada R, Satomi Y, Kurata T, Ueno N, Norioka S, Kondoh H, et al. Monounsaturated Fatty Acid Modification of Wnt Protein: Its Role in Wnt Secretion. Developmental Cell. 2006;11(6):791–801. doi: https://doi.org/10.1016/j.devcel.2006.10.003.

48. Rocks O, Peyker A, Kahms M, Verveer PJ, Koerner C, Lumbierres M, et al. An Acylation Cycle Regulates Localization and Activity of Palmitoylated Ras Isoforms. Science. 2005;307(5716):1746. doi: 10.1126/science.1105654.

49. Diez-Ardanuy C, Greaves J, Munro KR, Tomkinson NCO, Chamberlain LH. A cluster of palmitoylated cysteines are essential for aggregation of cysteine-string protein mutants that cause neuronal ceroid lipofuscinosis. Scientific reports. 2017;7(1):10-. doi: 10.1038/s41598-017-00036-8. PubMed PMID: 28127059.

50. Ayhan F, Perez BA, Shorrock HK, Zu T, Banez-Coronel M, Reid T, et al. SCA8 RAN polySer protein preferentially accumulates in white matter regions and is regulated by eIF3F. The EMBO Journal. 2018;37(19):e99023. doi: 10.15252/embj.201899023.

51. Zu T, Liu Y, Bañez-Coronel M, Reid T, Pletnikova O, Lewis J, et al. RAN proteins and RNA foci from antisense transcripts in C9ORF72 ALS and frontotemporal dementia. Proc Natl Acad Sci U S A. 2013;110:E4968–77. doi: 10.1073/pnas.1315438110. PubMed PMID: 24248382.

52. Albrecht AN, Kornak U, Böddrich A, Süring K, Robinson PN, Stiege AC, et al. A molecular pathogenesis for transcription factor associated poly-alanine tract expansions. Human Molecular Genetics. 2004;13(20):2351–9. doi: 10.1093/hmg/ddh277.

53. Oma Y, Kino Y, Toriumi K, Sasagawa N, Ishiura S. Interactions between homopolymeric amino acids (HPAAs). Protein Sci. 2007;16(10):2195–204. doi: 10.1110/ps.072955307. PubMed PMID: 17766374.

54. Kumar AS, Sowpati DT, Mishra RK. Single Amino Acid Repeats in the Proteome World: Structural, Functional, and Evolutionary Insights. PLOS ONE. 2016;11(11):e0166854. doi: 10.1371/journal.pone.0166854.

55. Pelassa I, Fiumara F. Differential Occurrence of Interactions and Interaction Domains in Proteins Containing Homopolymeric Amino Acid Repeats. Frontiers in Genetics. 2015;6(345). doi: 10.3389/fgene.2015.00345.

56. Green KM, Glineburg MR, Kearse MG, Flores BN, Linsalata AE, Fedak SJ, et al. RAN translation at C9orf72-associated repeat expansions is selectively enhanced by the integrated stress response. Nature Communications. 2017;8(1):2005. doi: 10.1038/s41467-017-02200-0.

57. Segev Y, Michaelson DM, Rosenblum K. ApoE ε4 is associated with eIF2α phosphorylation and impaired learning in young mice. Neurobiology of Aging. 2013;34(3):863–72. doi: https://doi.org/10.1016/j.neurobiolaging.2012.06.020.

58. Dorsman JC, Pepers B, Langenberg D, Kerkdijk H, Ijszenga M, den Dunnen JT, et al. Strong aggregation and increased toxicity of polyleucine over polyglutamine stretches in mammalian cells. Human Molecular Genetics. 2002;11:1487–96. doi: 10.1093/hmg/11.13.1487.

59. Oma Y, Kino Y, Sasagawa N, Ishiura S. Comparative analysis of the cytotoxicity of homopolymeric amino acids. Biochimica et Biophysica Acta (BBA) - Proteins and Proteomics. 2005;1748(2):174–9. doi: https://doi.org/10.1016/j.bbapap.2004.12.017.

60. van Eyk CL, McLeod CJ, O’Keefe LV, Richards RI. Comparative toxicity of polyglutamine, polyalanine and polyleucine tracts in Drosophila models of expanded repeat disease. Human Molecular Genetics. 2011;21(3):536–47. doi: 10.1093/hmg/ddr487.

61. Gurezka R, Laage R, Brosig B, Langosch D. A Heptad Motif of Leucine Residues Found in Membrane Proteins Can Drive Self-assembly of Artificial Transmembrane Segments. The Journal of Biological Chemistry. 1999;274(14):9265–70.

62. Whitley P, Grahn E, Kutay U, Rapoport TA, von Heijne G. A 12-Residue-long Polyleucine Tail Is Sufficient to Anchor Synaptobrevin to the Endoplasmic Reticulum Membrane. The Journal of Biological Chemistry. 1996;271(13):7583–6.

63. Cascarina SM, Ross ED. Proteome-scale relationships between local amino acid composition and protein fates and functions. PLoS computational biology. 2018;14(9):e1006256–e. doi: 10.1371/journal.pcbi.1006256. PubMed PMID: 30248088.

64. Rolls MM, Hall DH, Victor M, Stelzer EHK, Rapoport TA. Targeting of Rough Endoplasmic Reticulum Membrane Proteins and Ribosomes in Invertebrate Neurons. Molecular Biology of the Cell. 2002;13(5):1778–91. doi: 10.1091/mbc.01-10-0514.

65. Scior A, Preissler S, Koch M, Deuerling E. Directed PCR-free engineering of highly repetitive DNA sequences. BMC biotechnology. 2011;11:87. doi: 10.1186/1472-6750-11-87. PubMed PMID: 21943395; PubMed Central PMCID: PMC3187725.

66. Yanik MF, Cinar H, Cinar HN, Chisholm AD, Jin Y, Ben-Yakar A. Neurosurgery: functional regeneration after laser axotomy. Nature. 2004;432(7019):822. Epub 2004/12/17. doi: 10.1038/432822a. PubMed PMID: 15602545.

67. Schindelin J, Arganda-Carreras I, Frise E, Kaynig V, Longair M, Pietzsch T, et al. Fiji: an open-source platform for biological-image analysis. Nature methods. 2012;9(7):676–82. Epub 2012/06/30. doi: 10.1038/nmeth.2019. PubMed PMID: 22743772; PubMed Central PMCID: PMCPMC3855844.

68. Parslow A, Cardona A, Bryson-Richardson RJ. Sample drift correction following 4D confocal time-lapse imaging. Journal of visualized experiments : JoVE. 2014;(86). Epub 2014/04/22. doi: 10.3791/51086. PubMed PMID: 24747942; PubMed Central PMCID: PMCPMC4166950.

69. Yang JS, Nam HJ, Seo M, Han SK, Choi Y, Nam HG, et al. OASIS: online application for the survival analysis of lifespan assays performed in aging research. PloS one. 2011;6(8):e23525. doi: 10.1371/journal.pone.0023525. PubMed PMID: 21858155; PubMed Central PMCID: PMC3156233.

